# Influenza virus membrane fusion is promoted by the endosome-resident phospholipid bis(monoacylglycero)phosphate

**DOI:** 10.1101/2022.09.16.508338

**Authors:** Steinar Mannsverk, Ana M. Villamil Giraldo, Peter M. Kasson

## Abstract

The phospholipid bis(monoacylglycero)phosphate (BMP) is enriched in late endosomal and endo-lysosomal membranes and is believed to be involved in membrane deformation and generation of intralumenal vesicles within late endosomes. Previous studies have demonstrated that BMP promotes membrane fusion of several enveloped viruses, but a limited effect has been found on influenza virus. Here, we report the use of single-virus fusion assays to dissect BMP’s effect on influenza virus fusion in greater depth. In agreement with prior reports, we found that hemifusion kinetics and efficiency were unaffected by the addition of 10-20 mol % BMP to the target membrane. However, using an assay for fusion pore formation and genome exposure, we found full fusion efficiency to be substantially enhanced by the addition of 10-20 mol % BMP to the target membrane, while the kinetics remained unaffected. By comparing BMP to other negatively charged phospholipids, we found the effect on fusion efficiency mainly attributable to headgroup charge, although we also hypothesize a role for BMP’s unusual chemical structure. Our results suggest that BMP function as a permissive factor for a wider range of viruses than previously reported. We hypothesize that BMP may be a general co-factor for endosomal entry of enveloped viruses.

## Introduction

A shared feature of many enveloped viruses is that they subvert the endosomal pathway to gain entry into the host cell cytoplasm ^1^. The most common signal for triggering viral fusion is the gradually more acidic pH of the endosomal pathway, allowing the virus to time fusion to its preferred site in the endosome ^2^. In some cases, an additional receptor protein or protease present in the endosome is required for efficient fusion ^2,3^. However, although the role of pH and cellular proteins in viral entry is relatively well described, the role of compartment-specific lipids in viral entry is much less well understood ^4,5^.

bis(monoacylglycero)phosphate (BMP), formerly referred to as lysobisphosphatidic acid (LBPA), is a phospholipid that has only been detected in the endosomal and endolysosomal membrane of the cell, where it accounts for roughly 15-20 mol % of the total phospholipid content ^6–8^. The phospholipid has an unusual chemical structure, with each fatty acid tail being attached to separate glycerol moieties, which are in turn attached to a single phosphate group (**Fig. 3a**). BMP and its partner protein Alix have been shown to promote membrane deformation, regulate the biogenesis of intralumenal vesicles in late endosomes, control the fate of cholesterol and stimulate sphingolipid degradation ^9–11^. Interestingly, the presence of BMP in a lipid bilayer has shown to promote fusion of several enveloped viruses, including vesicular stomatitis virus (VSV), flaviviruses, phleboviruses and Lassa virus ^3,5,12–15^. However, a limited effect on influenza virus fusion has been found thus far ^3,13^.

Influenza A virus (hereafter referred to as influenza virus) is an enveloped virus belonging to the *Orthomyxoviridae* family, with a segmented, single-stranded, negative-sense RNA genome. The virus binds to terminal sialic acid residues on the host cell surface via its receptor binding protein, hemagglutinin (HA), and is subsequently endocytosed. Endosomal acidification triggers a conformational change in HA, exposing its fusion peptide and triggering fusion with mid-late endosomal membranes. Fusion proceeds through a hemifusion intermediate, where the proximal membrane leaflets mix while the distal leaflets remain separated, before a fusion pore is generated and expanded, facilitating the release of viral RNA segments into the cell cytoplasm ^16^.

In the past decade, single-virus fusion experiments have enabled the measurement of viral entry kinetics and efficiency ^17,18^ using infectious virus and either synthetically generated ^19^ or cell-derived membranes ^20,21^. When performed in microfluidic flow cells as opposed to live-cell tracking, these experiments permit precise control of the triggers for fusion as well as more facile manipulation of the membrane environment. Fluorescent reporters provide information on viral state changes, typically lipid mixing that is indicative of hemifusion and content release or genome exposure that is indicative of fusion pore formation ^17,18,22,23^.

Here, we leverage such single-virus fusion experiments to test the role of BMP in influenza virus membrane fusion, specifically examining hemifusion and fusion pore formation. We also compare BMP against other negatively charged phospholipids to understand the chemical basis for its effects. Our study helps shed light on how the endosomal lipid composition can modulate the complex replication cycle of influenza virus and how such lipids can act as general co-factors for enveloped viral fusion.

## Methods

### Materials

Palmitoyloleoylphophatidylcholine (POPC), dioleoylphosphatidylethanolamine (DOPE), cholesterol (CHOL), bis(monooleoylglycero)phosphate (S,R Isomer) (BMP), 1,2-dioleoyl-snglycero-3-phospho-(1’-rac-glycerol) (DOPG), 1,2-dioleoyl-sn-glycero-3-phospho-L-serine (DOPS) and biotinylated 1,2-dipalmitoyl-sn-glycero-3-phosphoethanolamine (biotin-DPPE) were acquired from Avanti Polar Lipids. Bovine brain disialoganglioside GD1a (Cer-Glc-Gal(NeuAc)-GalNAc-Gal-NeuAc) was purchased from Sigma-Aldrich. DiYO-1 (CAS 143413-85-8) was purchased from AAT Bioquest. Texas Red 1,2-dihexadecanoyl-sn-glycero-3-phosphoethanloamine (TR-DHPE) was purchased from Thermo Fisher. PLL-PEG and PLL-PEG-Biotin were purchased from SuSoS AG. X-31 influenza virus (A/Aichi/68, H3N2) was purchased at a titer of 6.3 × 10^9^ infectious units/mL from Charles River Laboratories. Reaction buffer consisted of 10 mM NaH_2_PO_4_, 90 mM sodium citrate and 150 mM NaCl.

### Liposome preparation and viral labelling

Liposomes were prepared as described elsewhere ^24^. In brief, the dried lipid film was hydrated in pH 7.4 reaction buffer containing 10µM DiYO-1, and large unilamellar vesicles with a nominal diameter of 100nm were generated by extrusion. **Table 1** lists the lipid composition of all liposomes used. The influenza virus envelope was fluorescently labelled with Texas Red-DHPE at a quenching concentration, as described elsewhere ^25^.

**Table 1.** Lipid composition of liposomes used. Percentage (%) signifies the mol % composition of the liposome.

| liposome name | POPC (%) | DOPE (%) | CHOL (%) | GD1a (%) | DPPE-Biotin (%) | additional lipid (%) |
| --- | --- | --- | --- | --- | --- | --- |
| 0 % BMP liposome | 57 | 20 | 20 | 2 | 1 | - |
| 10 % BMP liposome | 47 | 20 | 20 | 2 | 1 | BMP 10 |
| 20 % BMP liposome | 37 | 20 | 20 | 2 | 1 | BMP 20 |
| 20 % DOPG liposome | 37 | 20 | 20 | 2 | 1 | DOPG 20 |
| 20 % DOPS liposome | 37 | 20 | 20 | 2 | 1 | DOPS 20 |

### Electron cryomicroscopy

Sample vitrification was carried out using a Mark IV Vitrobot (ThermoFisher), according to the manufacturer’s instructions. 3µL of extruded liposomes were loaded onto a Quantifoil R 2/2 200 gold mesh carbon film grid, followed by a 5 min incubation and a 3 sec blotting step. Next, additional 3µL of sample was applied to the grid, followed by a 15 sec incubation and a 3 sec blotting step, before the grid was plunged into precooled liquid ethane. The long incubation time and double application were as recommended for liposome sample preparation ^26^. Sample screening and data acquisition were carried out on a 200kV Glacios electron microscope mounted with a Falcon III direct electron detector (ThermoFisher). Images were analysed using Fiji (version 2.3.0).

### Nanoparticle tracking analysis

Extruded liposomes were diluted 1:2000 in additional pH 7.4 reaction buffer and loaded into a NanoSight LM14 Nanoparticle Tracking Analysis Microscope (Malvern Panalytical) with a sCMOS camera attached, according to the manufacturer’s instructions. Analysis was carried out using the accompanying NanoSight NTA software (version 3.4). For each condition, an average size distribution plot was generated from three technical repeats consisting of 30 sec measurements.

### Single-virus fusion assays

Lipid mixing ^25^ and content mixing ^24^ of influenza particles with liposomes were performed as previously described. Liposomes decorated with DPPE-biotin were immobilised on the glass surface of a microfluidic flow cell via streptavidin linkage to PLL-PEG-biotin, displayed on an otherwise passivated surface. Next, virus was allowed to bind to the GD1a receptors displayed on the liposomes. Excess unbound virus was removed through buffer exchange before fusion was triggered by a rapid buffer exchange to pH 5 inside the flow cell chamber. All single-virus fusion assays were performed at 37 °C.

### Fluorescence microscopy, image analysis and statistics

Lipid mixing and genome exposure events were recorded via fluorescence video microscopy using a Zeiss Axio Observer inverted microscope with a 100x oil immersion objective and an sCMOS camera. Illumination and image acquisition were controlled using µManager ^27^. The microscope configuration and image acquisition parameters were identical to those recently described ^24^. Recorded images were analysed in MATLAB (The Mathworks, version R2021b), using previously developed single-virus detection and spot tracking code ^25,28^. The MATLAB code is available from https://github.com/kassonlab/micrograph-spot-analysis. All statistical tests were performed in MATLAB. Statistical tests for normal distribution of data and equal variance between conditions were performed using a Shapiro-Wilk normality test and multiple-sample test for equal variances, respectively.

## Results

### Lipid and content mixing between viral particles and liposomes containing BMP

Prior work on BMP suggested that it had a minimal effect on influenza hemifusion ^13^ or cell-cell fusion ^3^, but the effects on viral fusion kinetics and fusion pore formation had not been directly assessed. Here, we employed single-virus fusion measurements using both lipid mixing and a recently described content mixing assay ^24^ to test the effect of BMP on hemifusion and fusion pore formation.

Single-virus fusion experiments were performed using X-31 influenza virus bound to synthetic liposomes in microfluidic flow cells, as previously described ^25^. Target liposomes were generated to mimic the lipid composition of the late endosomal compartment ^8^, which included 10-20 mol % BMP (**Table 1**), and contained GD1a model glycosphingolipid receptors. After viral binding, fusion was triggered by a rapid buffer exchange and consequent drop in pH. Hemifusion was measured by Texas Red fluorescence dequenching upon viral particle lipid mixing with the target membrane. Fusion pore formation, also denoted full fusion, was measured by DiYO-1 fluorescence increase upon exposure of the viral interior to liposome contents, which permits the DiYO-1 dye to bind viral RNA. This binding is associated with a >100-fold increase in fluorescence quantum yield.

### Lipid mixing kinetics

Similar to prior reports for influenza ^28^, lipid mixing occurred rapidly after pH drop. The median time to lipid mixing was <7 seconds, and approximately 40% of labelled viral particles underwent lipid mixing within 5 min (**Fig. 1a, 1b**). Moreover, the addition of 10-20 mol % BMP to the target membrane did not alter the kinetics or efficiency of lipid mixing (**Fig. 1a, 1b**), in accordance with previous reports ^13^.

**Figure 1.**
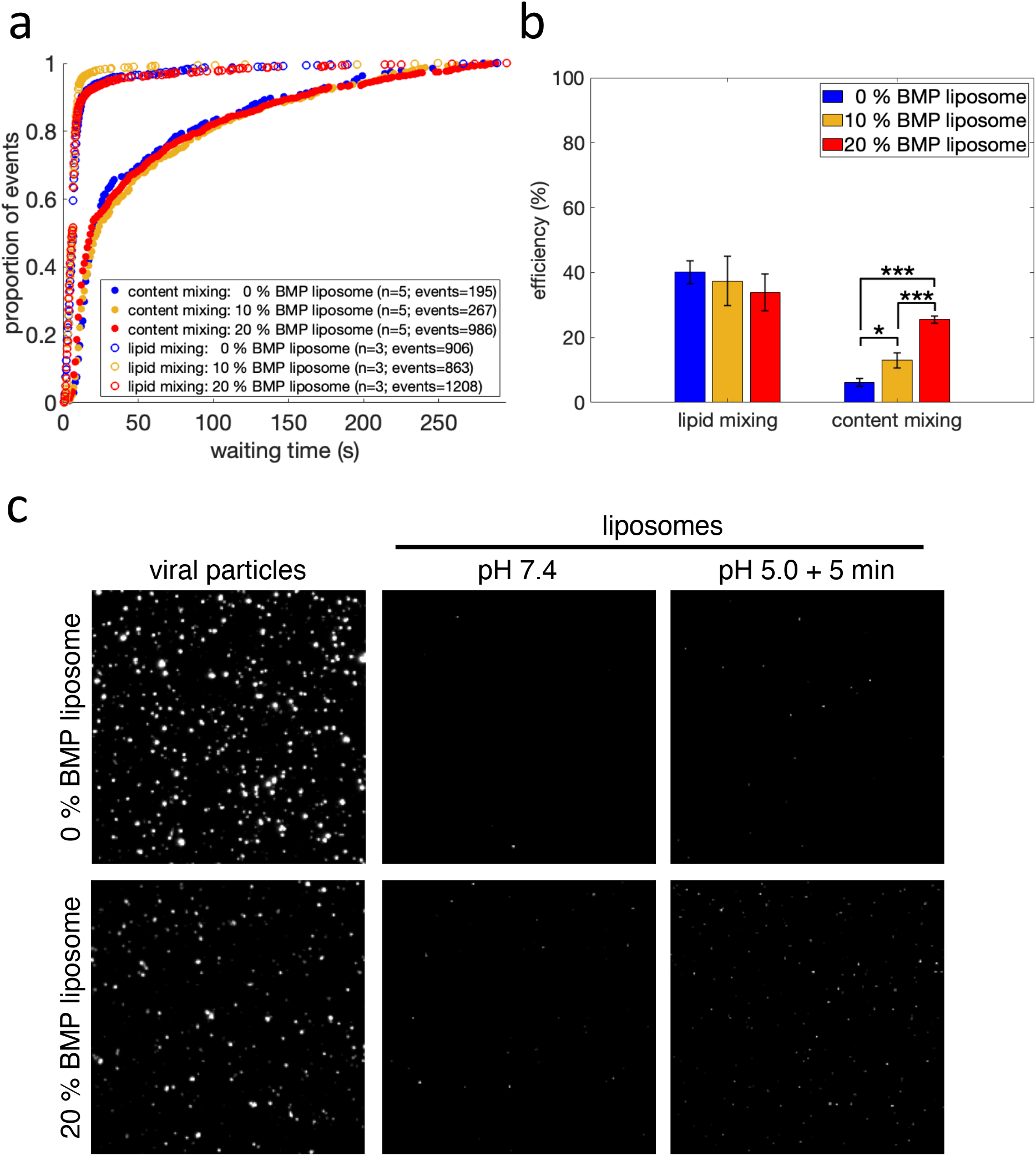
influenza virus lipid and content mixing with liposomes containing BMP. Lipid and content mixing kinetics are plotted in (**a**) as cumulative distribution functions of mixing events versus time after pH drop. The number of independent flow cell channels used to acquire the data is denoted “n” and the total number of mixing events is denoted “events”. Efficiency, defined as the number of lipid or content mixing events between a viral particle and a liposome divided by the total number of labelled viral particles, is plotted in (**b**). Values are plotted as mean +/-standard error, * denotes p-value < 0.05, and *** denotes p-value < 0.001, as determined by a one-way ANOVA test (f-value 36.78; p-value = 7.61 × 10^−6^) and a Tukey-Kramer post-hoc test. The number of independent channels and events recorded are also displayed in (**a**). Fluorescence micrographs are rendered in (**c**), showing viral particles undergoing content mixing with target liposomes containing 0 mol % BMP (top row) or 20 mol % BMP (bottom row). “Viral particles” (first column) displays membrane-labelled viral particles and “liposomes” displays liposomes before (second column) and 5 min after the pH drop (third column). White spots visualized after but not before pH drop represent liposomes that have undergone content mixing with virus.

### Content mixing kinetics

Content mixing occurred more slowly than lipid mixing and with a lower efficiency (**Fig. 1a, 1b**), as expected for a later step in influenza membrane fusion. The presence of 10-20 mol % BMP in the target membrane did not affect the kinetics of content mixing (p-value > 0.6 via bootstrapped rank sum test; **Fig. 1a**). However, we observed a dose-dependent increase in content mixing efficiency, in the presence of 10-20 mol % BMP in the target membrane (**Fig. 1b**). In the absence of BMP, approximately 6% of labelled viral particles underwent content mixing, while 10 and 20 mol % BMP in the target membrane resulted in an approximately twofold and fourfold increase in content mixing efficiency, respectively (**Fig. 1b**). Sample micrographs of content mixing after pH drop are shown in **Fig. 1c**. A small number of liposomes were fluorescent in the DiYO-1 channel prior to pH drop (**Fig. 1c**). In the absence of virus, no liposomes were fluorescent in the DiYO-1 channel, suggesting that these represent rare interactions with some element of the viral sample at neutral pH. However, these fluorescent liposomes did not undergo further fluorescence increase after the pH drop and consequently did not contribute to the overall recorded content mixing events.

### Lipid morphology and size

To test whether the observed increase in full fusion efficiency could result from differences in the size or morphology of the liposomes containing BMP, we compared the morphology and size of our liposomes containing 0 and 20 mol % BMP in their membrane. Prior studies have noted that when resuspended, 100 mol % BMP forms non-spherical liposomes with bud-like surface protrusions and that extruded liposomes containing 100 mol % BMP have a smaller diameter than similar liposomes containing POPG or POPC lipids ^29^. Even if the composition range of BMP in our liposomes is ≤20 mol %, we wanted to rule this out.

Electron cryomicrographs showed that liposomes extruded at 100 nm containing 0 or 20 mol % BMP formed mostly unilamellar, spherical liposomes (**Fig. 2a**). Single-particle analysis of particle size via Brownian diffusion (nanoparticle tracking) yielded a size distribution with two major modes: ∼100nm and 140±6nm in diameter (**Fig. 2b**). The size distribution between the 0 and 20 mol % BMP liposome is comparable, with mean diameters of 128nm and 126nm, respectively (**Fig. 2b**). These results demonstrate that the observed differences in full fusion efficiency when adding BMP to the target liposomes are not attributable to changes in liposome morphology or size.

**Figure 2.**
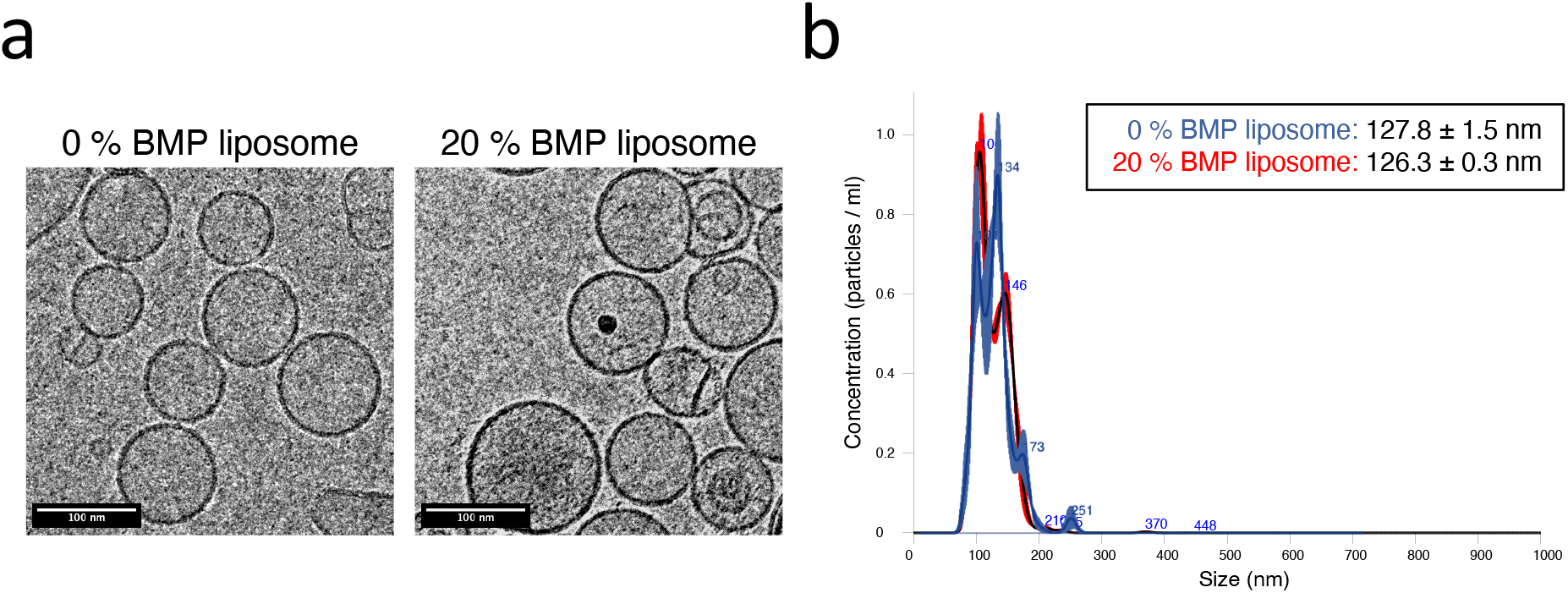
20 mol % BMP liposome morphology and size distribution. Representative electron cryomicrographs of liposomes containing 0 and 20 mol % BMP are rendered in (**a**), demonstrating that both liposomes have similar morphology after extrusion through a 100nm filter membrane. 16 micrographs were acquired in total; additional micrographs are displayed in **Fig. S2**. Liposome size distributions determined via nanoparticle tracking analysis are plotted in (**b**). Plotted distributions show the average of three technical repeats measured for 30 sec each. 0 mol % (blue) and 20 mol % (red) BMP liposomes were measured <6 hours after extrusion through a 100nm filter membrane. Legend lists mean liposome size +/-standard error. The measurement was repeated on three independent lipid extrusions, with equivalent size distributions observed.

### Influenza fusion to liposomes containing other negatively charged lipids

BMP has two distinctive features: a negatively charged headgroup and an *sn-1:sn-1’* glycerophosphate stereoconfiguration with a fatty acid attached to each of its glycerol moieties ^30^. To probe the basis for the observed effects of BMP, we replaced the 20 mol % BMP in the target membrane with a structural isomer and precursor of BMP, dioleoyl phosphatidylglycerol (DOPG) ^31,32^. DOPG has a slightly different chemical structure in that it possesses the more common *sn-3:sn-1’* stereoconfiguration and has both fatty acids attached to one glycerol moiety (**Fig. 3a**). Replacing 20 mol % BMP in the target membrane with DOPG did not affect hemifusion kinetics or efficiency. It did, however, result in a modest, non-significant (p = 0.38 via bootstrapped rank sum test) slowing of full fusion kinetics (**Fig. 3b, 3c**). Interestingly, DOPG and BMP exhibited identical full fusion efficiencies (**Fig. 3c**).

**Figure 3.**
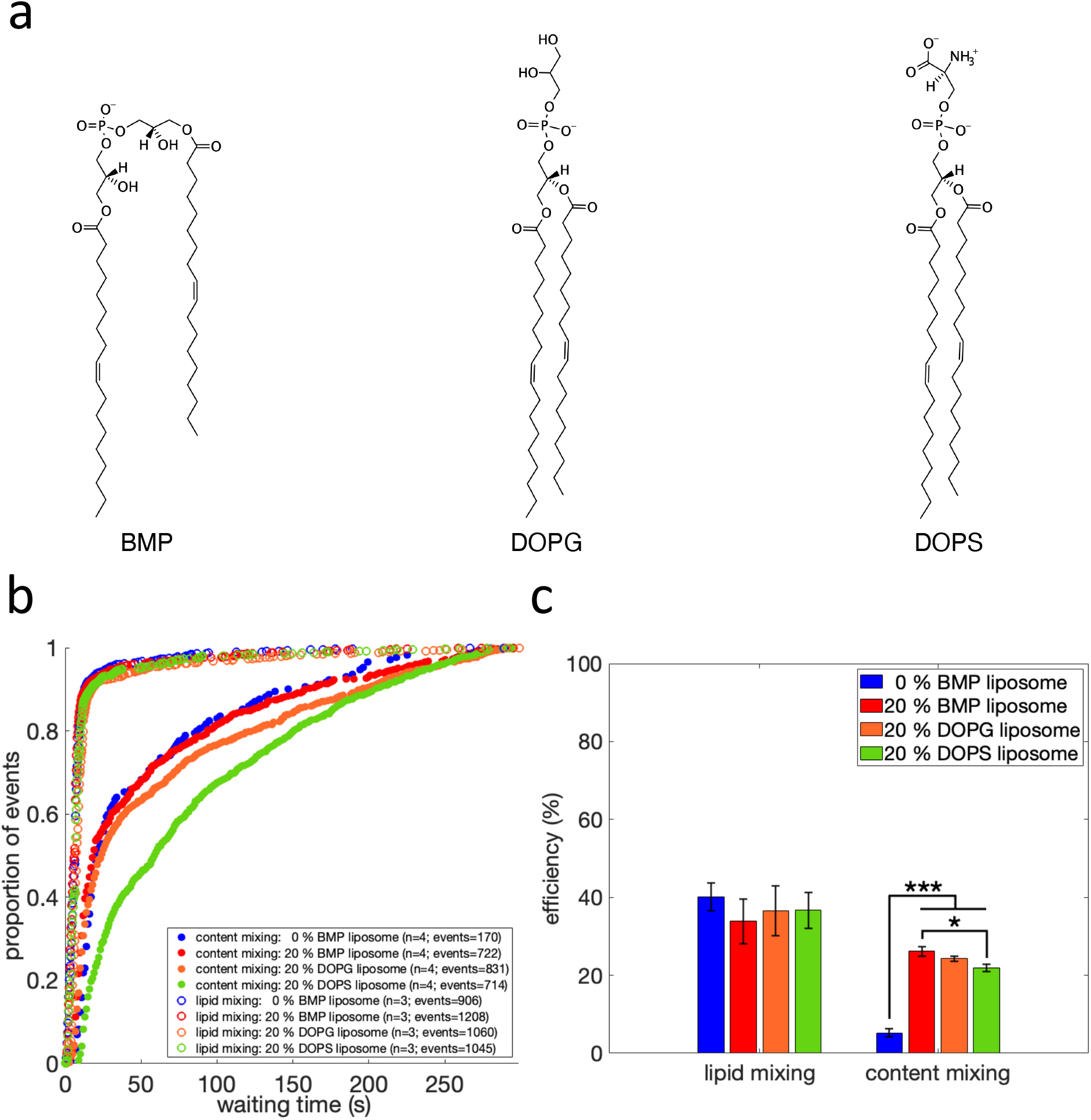
Chemical structure, lipid and content mixing of liposomes containing anionic phospholipids. Rendered in (**a**) are the chemical structures of BMP (left), DOPG (middle) and DOPS (right). Lipid and content mixing kinetics are plotted in (**b**) as cumulative distribution functions of mixing events versus time after pH drop, for liposomes containing 0 mol % BMP or 20 mol % BMP, DOPG or DOPS. The number of independent flow cell channels used to acquire the data is denoted “n” and the total number of mixing events is denoted “events”. Lipid and content mixing efficiencies are plotted in (**c**). Values are plotted as mean +/-standard error, * denotes p-value < 0.05 and *** denotes p-value < 0.001, as determined by a one-way ANOVA test (f-value = 92.38; p-value = 1.47 × 10^−8^) and a Tukey-Kramer post-hoc test. Number of independent channels and events recorded are displayed in (**b**). Cumulative distribution functions for content mixing are plotted with bootstrapped confidence intervals in **Fig. S3**.

Since anionic phospholipids have been shown to promote fusion of enveloped viruses ^5,12^, we also replaced BMP in the target membrane with a structurally less-related anionic phospholipid, dioleoyl phosphatidylserine (DOPS) (**Fig. 3a**). 20 mol % DOPS in the target membrane did not affect hemifusion kinetics or efficiency, but we observed a slowing of full fusion kinetics (p < 0.001 via bootstrapped rank sum test; **Fig. 3b, 3c**). Moreover, we observed a very modest decrease in full fusion efficiency compared to BMP (**Fig. 3c**).

Since the presence of both the anionic phospholipids DOPG and DOPS in the target membrane also resulted in an increase in pore formation efficiency during influenza virus fusion (**Fig. 3c**), we believe that the negatively charged headgroup of BMP contributes substantially to the observed increase in full fusion efficiency. However, DOPG and DOPS both displayed slower full fusion kinetics than BMP (**Fig. 3b**), suggesting that the unusual chemical structure of BMP also plays an important role during influenza virus fusion. Example fluorescence micrographs of content mixing events between viral particles and 20 mol % BMP, DOPG or DOPS liposomes are shown in **Fig S1**. It should be noted that the BMP, DOPG and DOPS molecules used all have identical acyl tail composition (18:1, Δ9-Cis), hence the observed differences are not attributable to this.

## Discussion

Previous studies have shown that the presence of the phospholipid BMP in the target membrane promotes hemifusion of several enveloped viruses ^12–15^. These studies did not find an effect for BMP on influenza hemifusion, and our results support that conclusion. Here we demonstrate that the presence of BMP in the target membrane during influenza virus fusion greatly enhances the likelihood that a hemifusion intermediate progresses to form a fusion pore. This finding is similar to results on other enveloped viruses ^3,5,12,15^.

Potential explanations for the effect of BMP in promoting fusion pore formation include 1) specific interactions with fusion peptides or fusion loops, 2) change in spontaneous negative curvature of the target membrane or 3) an effect on fusion pore opening specific to the chemical structure of BMP. Prior studies on BMP and dengue or VSV postulated that lipid-peptide interactions may be responsible for some of the effects observed ^5,12^. It is possible that such effects also exist for influenza, although as BMP is implicated in the entry of a greater number of viral families a specific peptide interaction becomes less likely. Nonetheless, it is possible that a more general phenomenon, such as charge-charge interactions between the peptides and the distal membrane leaflet, could be responsible for promoting progression past the hemifusion stage ^33^.

Negative spontaneous curvature has been employed as a unifying concept for understanding the effect of several lipids on promoting viral membrane fusion ^3,15,34–38^. Both our study on influenza and prior work on other enveloped viruses found multiple anionic lipids capable of promoting fusion ^5,12,15^, and all of these lipids exert a negative spontaneous curvature on lipid bilayers ^39,40^. Therefore, the bulk membrane energetics of BMP-containing membranes may be important in stabilizing high-energy fusion intermediates and promoting progression to full fusion. Similar effects have been found for cholesterol ^19^, which has multiple activities in membranes but also promotes negative spontaneous curvature ^41^.

In addition to spontaneous curvature, negative charge is a common chemical feature of many fusion-promoting lipids. Interestingly, VSV showed a specific preference for BMP over phosphatidylserine, while dengue virus lipid mixing was promoted by several anionic lipids ^5,13^. However, BMP is likely the anionic lipid most relevant for endosomal entry of viruses, as phosphatidylglycerol has not been detected and phosphatidylserine accounts for less than 3% of the total phospholipid content of the late endosomal membrane ^8^, where influenza virus fusion occurs. Our work on influenza and studies on other enveloped viruses have found differences in fusion kinetics between BMP and phosphatidylserine or phosphatidylglycerol ^12,13^. It is thus possible that the unusual chemical structure of BMP plays an additional role in promoting fusion, perhaps stabilizing key intermediates. BMP is specifically enriched in highly curved multivesicular bodies within endosomes and is believed critical to their stability ^7,9^, so it is possible that this helps explain the endosomal entry preference of many enveloped viruses.

For BMP to drive intralumenal vesicle formation in late endosomes, a proton gradient must exist across the membrane ^9^. Coincidentally, in our experimental setup the same proton gradient exists during influenza virus fusion with target liposomes: upon fusion triggering, the viral particle is located in a lumen-like environment (pH 5.0) and fusion occurs towards the liposome interior, which exhibits a cytoplasmic-like environment (pH 7.4). It would be interesting to investigate whether this proton gradient is essential for BMP’s effect on influenza virus fusion.

## Conclusions

Our findings show that BMP plays an important role in promoting fusion pore formation during influenza virus fusion. BMP alters fusion pore formation efficiency rather than kinetics, suggesting that it likely modulates flux between alternative kinetic pathways rather than simply altering a free-energy barrier in a committed process. This endosomally enriched phospholipid has now been shown to enhance entry in multiple viral families that enter via the endosomal compartment. We speculate that it thus exhibits a general mechanism of promoting viral membrane fusion. Furthermore, the presence of BMP in the endosomal membrane may partly explain why so many enveloped viruses enter via the endosomal pathway rather than the plasma membrane, despite the risk of proteolytic degradation that this pathway entails.

## Supporting Information

Fluorescence micrographs of additional content-mixing events, additional electron cryomicrographs, and uncertainty analysis of content-mixing kinetics (PDF).

## Acknowledgements

We acknowledge the use of the Cryo-EM Uppsala facility for sample preparation and data acquisition. The authors thank G. Melikian for many helpful conversations. This work was supported by a Wallenberg Academy Fellowship to P.M.K. and by the Swedish Research Council (2017-04236 to P.M.K.).

