## Supporting Information for "Influenza virus membrane fusion is promoted by the endosome-resident phospholipid bis(monoacylglycero)phosphate"

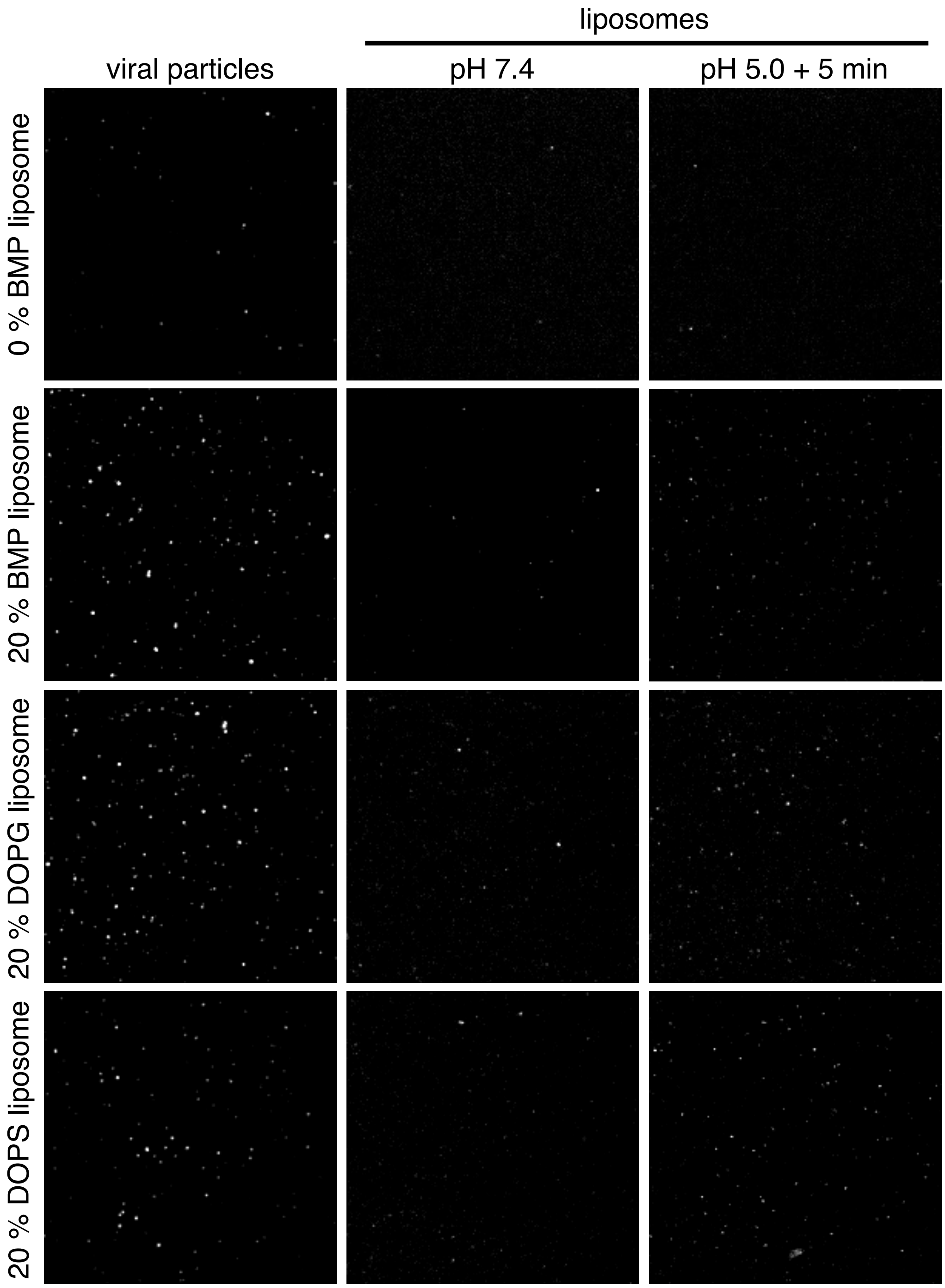


**Supporting Figure 1. Additional micrographs of content mixing events with liposomes containing anionic phospholipids.** Fluorescence micrographs shows viral particles undergoing content mixing with target liposomes containing 0 mol % BMP (first row), 20 mol % BMP (second row), 20 mol % DOPG (third row) or 20 mol % DOPS (fourth row). “Viral particles” (first column) displays membrane-labelled viral particles and “liposomes” displays liposomes before (second column) and 5 min after the pH drop (third column). White spots visualized after but not before pH drop represent liposomes that have undergone content mixing with virus. A small number of liposomes displayed DiYO-1 fluorescence prior to the pH drop (second column).

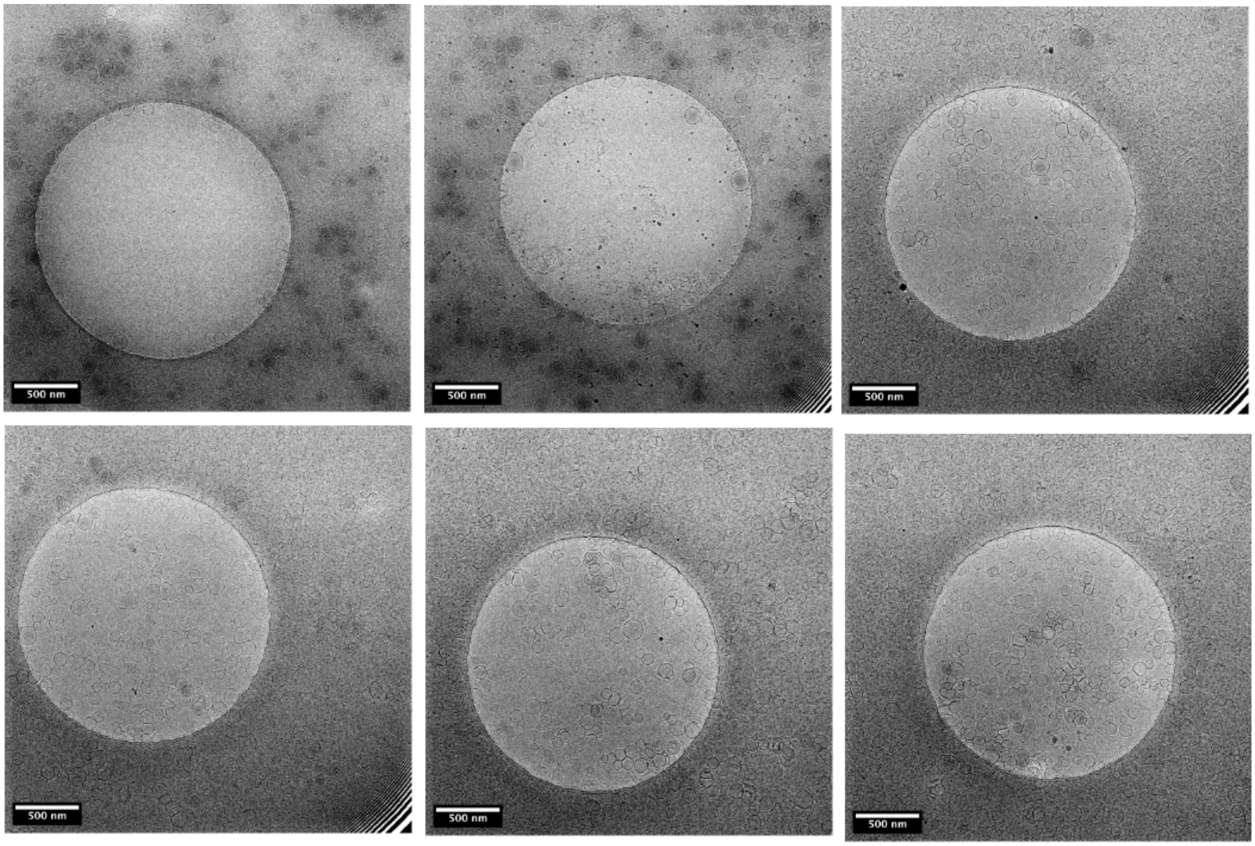


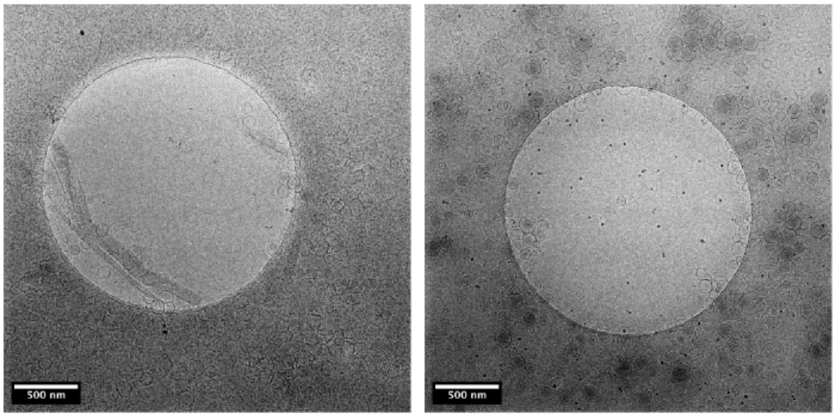


0% BMP liposomes

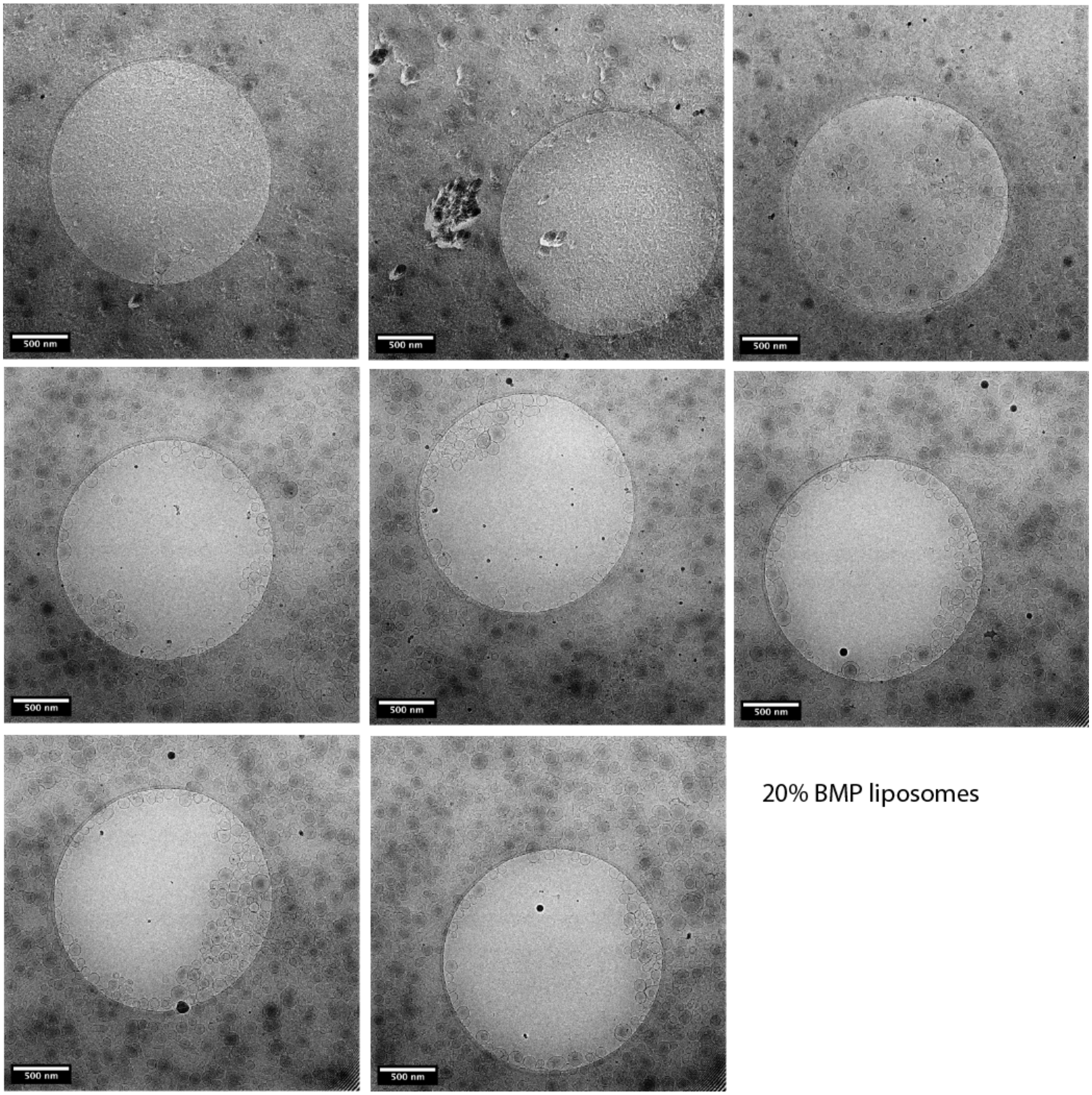


**Supporting Figure 2. Images of all electron micrographs acquired**. Thumbnail images of electron micrographs acquired are displayed, 8 at 0% BMP and 8 and 20% BMP.

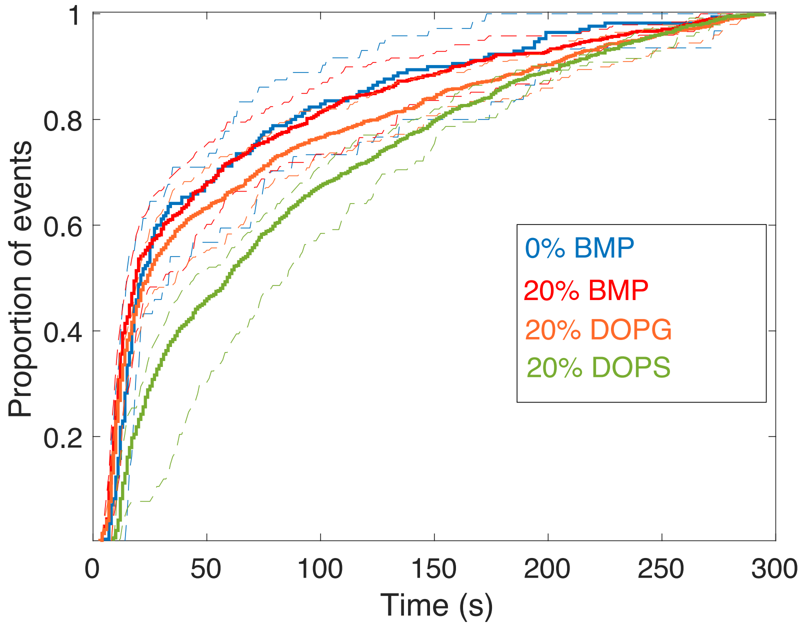


**Supporting Figure 3. Uncertainty analysis of content-mixing kinetics**. Uncertainty distributions were quantified for content-mixing experiments via bootstrap resampling across flow cell channels (experimental replicates). Cumulative distributions are plotted with the observed CDF in solid lines and the 90% confidence intervals in dashed lines. Distributions were assessed as different or not via bootstrapped rank sum tests of individual-virus waiting times, bootstrap resampled across flow cell channels.
